# Identification of key residues of the DNA glycosylase OGG1 controlling efficient DNA scanning and recruitment to oxidized bases in living cells

**DOI:** 10.1101/2022.11.04.515179

**Authors:** Ostiane D’Augustin, Virginie Gaudon, Capucine Siberchicot, Rebecca Smith, Catherine Chapuis, Jordane Depagne, Xavier Veaute, Didier Busso, Anne-Marie Di Guilmi, Bertrand Castaing, J. Pablo Radicella, Anna Campalans, Sébastien Huet

**Author notes:** Corresponding authors: Anna Campalans and Sébastien Huet.

## Abstract

The DNA-glycosylase OGG1 oversees the detection and clearance of the 7,8-dihydro-8-oxoguanine (8-oxoG), which is the most frequent form of oxidized base in the genome. This lesion is deeply buried within the double-helix and its detection requires careful inspection of the bases by OGG1 via a mechanism that remains only partially understood. By analyzing OGG1 dynamics in the nucleus of living human cells, we demonstrate that the glycosylase constantly scans the DNA by rapidly alternating between diffusion within the nucleoplasm and short transits on the DNA. This scanning process, that we find to be tightly regulated by the conserved residue G245, is crucial for the rapid recruitment of OGG1 at oxidative lesions induced by laser micro-irradiation. Furthermore, we show that residues Y203, N149 and N150, while being all involved in early stages of 8-oxoG probing by OGG1 based on previous structural data, differentially regulate the scanning of the DNA and recruitment to oxidative lesions.

## INTRODUCTION

DNA bases can be oxidized by reactive oxygen species arising from intracellular metabolism or external stresses. Due to its low oxidative potential (1), guanine is the most frequent target for oxidation among the different bases, leading to the formation of 7,8-dihydro-8-oxoguanine (8-oxoG). This base modification, which has been associated with various cancers and age-related diseases (2– 5), promotes G:C to T:A transversions due to its ability to pair with adenine through its s*yn* conformation (6). It is therefore crucial to efficiently clear 8-oxoG from the genome to avoid accumulation of mutations. In mammals, 8-oxoG facing a cytosine is recognized and excised by the 8-oxoguanine DNA-glycosylase 1 (OGG1) (7–10), hence initiating the base excision repair (BER) pathway (11, 12).

Unlike many other DNA lesions, 8-oxoG does not induce helix distortion nor blockage of either transcription or replication (13–16). Therefore, to efficiently detect and excise 8-oxoG, OGG1 needs to constantly scan the DNA through mechanisms not yet completely characterized. Structural data have revealed different conformations of the OGG1/DNA complex that could correspond to consecutive steps of this base inspection process (17–19). The binding of OGG1 to DNA leads to a local kinking of the DNA (20), which in turn promotes the flipping-out of the base for further inspection and potential cleavage. While it is unclear whether extrusion occurs irrespectively of the oxidation state of guanine, recent data suggest that only 8-oxoG is stabilized in this extra-helical position which facilitates the entry of the modified base into the OGG1 catalytic pocket (19). Importantly, we still miss a full cartography of the OGG1 amino-acid residues involved in these early steps of 8-oxoG detection. Residues such as the Y203 or N149 (in all this work, the residue numbering refers to the human OGG1) were shown to be important for intrahelical lesion sensing or for specificity towards the 8-oxoG:C pair (17, 19), but other conserved residues localized further away from the extruded base could also participate to these probing steps.

Besides these elementary molecular steps of the recognition of 8-oxoG by OGG1, the mechanisms regulating the exploration of the nuclear environment by OGG1 to ensure rapid detection of the oxidized guanines remain unclear. *In vitro* data demonstrated that OGG1 is able to diffuse along the DNA while searching for 8-oxoG (21), these DNA scanning phases probably alternating with periods of free diffusion to travel from one genomic location to another (22). Although this search process is thought to be impacted by the wrapping of the DNA around histones (23), the dynamics of OGG1 within the nucleus remains poorly characterized (24).

In this work, we establish fluorescence-based assays to quantitatively characterize the dynamic behavior of OGG1 in the nucleus of living cells. Combining microscopy and biochemistry approaches, we were able to determine how conserved residues of OGG1 thought to be involved at early steps of lesion recognition impact the navigation of the glycosylase within the nucleus and its accumulation onto 8-oxoG induced by laser micro-irradiation.

## MATERIAL AND METHODS

### Plasmids

All plasmids used in this study are listed in Table S1. The following plasmids were previously described: APE1-GFP (25), GFP2 and GFP5 (26). All OGG1 plasmids refer to the α isoform of human OGG1. For all the fluorescently tagged OGG1 constructs, a linker (translation PDPSGAAAAGGSQK) was inserted between OGG1 and EGFP in the plasmid previously described (27). Briefly, original OGG1-GFP plasmid was amplified by PCR with Phusion DNA polymerase using primers bringing the linker sequence (Table S2) using previously described method (28). Upon amplification, the PCR product was treated overnight with DpnI at 37°C to digest the original plasmid, and transformed in DH5α-T1R home-made competent cells. Point mutations in OGG1-L1-GFP were generated using site-directed mutagenesis with primers listed in Table S2. To generate the GST-OGG1 fusion proteins, the wild-type and OGG1 (G245A) coding sequences were amplified by PCR with Phusion DNA polymerase (NEB) (primers indicated in Table S2) and PCR products were cloned in pnEAvG-based plasmids (29) by complementary single-strand annealing cloning (30). Upon amplification, PCR products were purified on GeneJet PCR purification kit (Thermo Fisher Scientific) to remove all dNTPs. The pnEAvG plasmid was digested by NdeI + BamHI. DNA molecules were treated with T4 DNA polymerase to generate complementary single strands. Upon annealing, the reaction was transformed in DH5α-T1R home-made competent cells. For the generation of the plasmids coding for the His-tagged OGG1, OGG1 sequence was PCR amplified from pPR71 (8) cloned into NdeI/EcoRI sites of the overexpression vector pET30a (Novagen). The recombinant pET30a-hOGG1 was used as a template to introduce Y203A and N149A/N150A (referred to as 2NA) substitutions using the Quick-Change kit (Agilent) using primers indicated in Table S2. All constructs were validated by DNA sequencing. Rescriction enzymes and DNA polymerases were from New England Biolabs, antibiotics from Sigma-Aldrich and primers from Eurogentec.

### Cell lines and treatments

The HeLa OGG1 knockout cell lines were generated according to the protocol described by the Zhang lab (31). The target sequence for *OGG1* was chosen according to the web-based CRISPR design tool CRISPick (https://portals.broadinstitute.org/gppx/crispick/public). The sgRNA oligos (Table S2) were introduced into pX458 expressing Cas9 nuclease fused to GFP (Addgene #48138). pSpCas9(BB)-2A-GFP (PX458) was a gift from Feng Zhang (Addgene plasmid #48138). HeLa WT cells were transfected with the guide containing plasmid using the transfection reagent XtremeGENE HP (Roche) according to manufacturer’s protocol. Single GFP positive cells were sorted into 96-well plates using the FACSAria Fusion flow cytometer (BD Biosciences). The knockout cell lines grown from the single cells were screened by western blot using an antibody against OGG1 (Abcam, ab124741). Cells were cultured in DMEM (Thermo Fisher Scientific) containing 10% fetal bovine serum (FBS) (Eurobio) and 1% of penicillin and streptomycin (PS) (Thermo Fisher Scientific) at 37°C with 5% CO_2_. Cells were plated in chambered coverglass (Zell Kontact or Thermo Fisher Scientific) and transfected with the indicated plasmid 48 to 72 hrs prior to imaging using X-Treme gene HP (Roche) transfectant reagent according to the manufacturer’s instructions. Immediately before imaging, growth medium was replaced with gas-balanced phenol red-free Leibovitz L-15 imaging medium (Thermo Fisher Scientific) supplemented with 20% FBS and 1% PS. TH5487 (Probechem, PC-35806) and O8-Cl (Tocris) were added at 30μM in the imaging medium 20 to 30 mins prior to imaging. Fresh imaging medium containing O8-Cl was renewed every 2 hrs due to the low stability of this compound.

### Immunostaining

To validate the induction of 8-oxoG by laser micro-irradiation, a dozen of cells was micro-irradiated and immediately fixed on ice with cold methanol and acetone (1:1), air dryed, and rehydrated in PBS. DNA was denatured with 2N HCl for 45 mins, neutralized with Tris-HCl pH 8.8 for 5 min and washed three times in PBS. Then, cells were permeabilized with PBS-Triton 0.1% and incubated in blocking buffer (BSA 3%, Triton 0.1% in PBS) for 1 hr. All these steps were performed at room temperature (RT). Cells were incubated at 4°C overnight with an anti-8-oxoG mouse antibody (Abcam, ab48508) used at 0.6 μg/mL in blocking buffer. After three consecutive washing in PBS-Triton 0.1% for 10 mins, cells were incubated at RT for 1 hr with an anti-mouse secondary antibody tagged with AlexaFluor488 (Thermo Fisher Scientific, A11001) used at 2 μg/mL in blocking buffer. DNA was stained for 10 min at RT with Hoechst in PBS (10μg/mL). Cells were rinsed 3 times and transferred in PBS for imaging. For the colocalization experiments, cells expressing unfused GFP2 or OGG1-GFP were fixed in 4% paraformaldehyde for 15 min at RT and DNA was stained with Hoechst as described above.

### Protein purification

To obtain protein extracts from WT and OGG1 KO cells, frozen cells (approx. 5×10^6^ cells) were resuspended in 100 μL of lysis buffer (25 mM Tris pH 7.5, 250 mM NaCl, 1 mM EDTA), sonicated for 10 s on ice (1s pulses with 10s intervals, using a Branson digital sonicator, power set to 10%) and centrifuged 90 mins at 14 000 rpm and 4°C. Protein concentrations of the supernatants were determined by the Bradford method (BioRad).

Plasmids pnEAvG-HsOGG1a and pnEAvGHsOGG1-G245A coding for the GST-OGG1 WT and GST-G245A proteins, respectively, were transformed into *E. coli* BL21(DE3). Bacteria were grown at 37°C to reach OD_600_ = 1 when 0.5mM IPTG was added and the cultures switched to 20°C for 18 h. Cells were harvested, suspended in lysis buffer (20mM Tris at pH 8 and 4°C, 500mM NaCl, 0.1% NP40, 2mM EDTA, 2mM DTT, 10% glycerol, 1 mg/mL Lysozyme, 1mM 4-(2-aminoethyl) benzenesulphonyl fluoride, 10mM Benzamidine, 2μM pepstatine, 2μM leupeptine) and disrupted by sonication. Extract was cleared by centrifugation at 186.000 x g for 1 h at 4°C and then incubated at 4°C with Glutathione Sepharose™ 4B resin (GE Healthcare) for 3h. The mixture was poured into an Econo-Column Chromatography Column (BIO-RAD) and the beads were washed with buffer A (20mM Tris at pH 8 and 4°C, 500mM NaCl, 2mM EDTA, 2mM DTT, 10% glycerol) followed by buffer B (20mM Tris at pH 8 and 4°C, 100mM NaCl, 2mM EDTA, 2mM DTT, 10% glycerol). GST fusions were eluted with Buffer B containing 30 mM glutathion. Fractions containing GST-protein were pooled and loaded to a 6 mL Resource S column (GE Healthcare) equilibrated with buffer B. The protein was eluted with a 90 mL linear gradient of 0.1–0.5 M NaCl. Peak fractions containing purified GST-OGG1 were pooled and glycerol was added to reach a final concentration of 50%. The His-tagged WT OGG1 and the mutated versions Y203A and 2NA were produced and purified as previously described (32). The 6-His-Tag of the protein of interest was eliminated by enterokinase digestion. The resulting proteolysate was applied onto a benzamidin column (GE Healthcare) to eliminate enterokinase and then onto a HisTrapTM (GE Healthcare) equilibrated in 20 mM Hepes/NaOH, pH 7.6, 500 mM NaCl, 5% glycerol, 5 mM Imidazole and 1 mM DTT. The untagged protein eluted in the flow-through was then loaded on POROSTM HS20 cation exchanger equilibrated in buffer C (20mM Hepes pH7.6, 5% glycerol, 1mM DTT) and eluted with a linear NaCl gradient with buffer D (= buffer C + 1M NaCl). Elution fractions containing the untagged protein were pooled and the homogenous protein was finally isolated by size-exclusion chromatography onto a Superdex 75 column (GE Healthcare) equilibrated in 20 mM Hepes/NaOH, pH 7.6, 350 mM NaCl, 5% glycerol and 1 mM TCEP, concentrated at about 200 μM and stored at -80°C.

### Electrophoretic mobility shift assay

The ability of the OGG1 variants to bind the oligonucleotides indicated in Table S2 was analyzed by electrophoretic mobility shift assay (EMSA). Cy5 labelled DNA duplexes containing a G:C or an 8-oxoG:C pair used at 5 nM were incubated with increasing concentrations of OGG1 (G245A vs WT) for 30 min at 4°C in a buffer containing 50 mM Tris-HCl pH 7.5, 115 mM NaCl, 1 mM EDTA, 0.06 mg/mL BSA, 0.6 mM DTT, 7.5% Glycerol. The other variants (Y203A and 2NA vs WT) were studied as described previously (33). Briefly, 0.1 nM of 5’-[^32^P]-labeled 24-mer DNA duplexes [G:C] or [8-oxoG:C] were incubated at 4°C for 30 min, alone or in the presence of the indicated protein concentrations, in 20 mM Hepes/NaOH, pH 7.6, 100 mM NaCl, 10% Glycerol, 0.1% BSA and 0.5 mM TCEP. Binding reactions were loaded onto a non-denaturing 10% polyacrylamide gel for electrophoresis as described previously (34). After electrophoresis (14V/cm at 4°C), dried gels were exposed for autoradiography using a Typhoon Molecular Imager (Amersham) and quantified using ImageQuant software. Triplicate EMSA titration experiments were performed to extract apparent dissociation constants K_Dapp_ which is close to the enzyme concentration needed for half-maximal binding under the experimental conditions chosen (34). We used the Origins software (version 9.0.0, OriginLab) to fit the curves of the relative amounts of DNA/protein complex (C) as a function of the protein concentration (P) with a non-linear regression logistics function according to Hill equation:

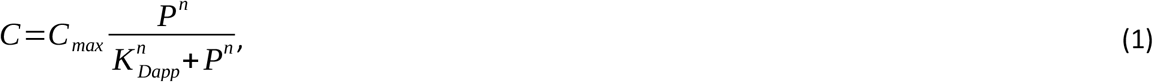

where *C*_*max*_ is the maximal amount of DNA/protein complex, *K*_*Dapp*_ is the apparent dissociation constant and *n* is the Hill coefficient.

### OGG1 DNA glycosylase activity

An oligonucleotide containing an 8-oxoG annealed with the complementary strand (Table S2) was used to evaluate OGG1 DNA glycosylase activity of HeLa WT or OGG1 KO protein extracts and proteins purified from *E. coli*. Total protein extracts or GST-tagged OGG1-WT and OGG1-G245A purified proteins, were incubated with 180 fmoles of the Cy5-labelled oligonucleotide at 37°C in a final volume of 30 μL of reaction buffer (25 mM Tris pH 7.5, 80 mM NaCl, 1 mM EDTA) for 15 mins or as indicated. Reactions were stopped by adding 2 μL of 1.5 M NaOH and further incubated for 15 mins at 37°C to cleave the remaining abasic sites. 4 μL of formamide loading buffer was added before heating 5 min at 95°C. The DNA glycosylase activity of the other purified mutants was measured using 20 nM of 5’-[^32^P]-labeled 24-mer [8-oxoG:C] incubated at 37°C for 15 mins, alone or in the presence of indicated concentrations of OGG1 (WT, 2NA or Y203A) in TBE (1X) containing 150 mM NaCL and 0.1% BSA. The reaction was stopped by 5 min incubation with 0.2 M NaOH at 50°C followed by 3 mins incubation with formamide dye loading buffer at 75°C. Substrate and products were separated by electrophoresis on a 20% denaturing polyacrylamide gel (urea-PAGE: 19:1 acrylamide/bisacrylamide in TBE1X and 7 M urea) for 30 mins at 400 V. Gel autoradiography was performed using a Typhoon Molecular Imager (Amersham) and quantified using ImageQuant software.

### Fluorescence imaging and correlation spectroscopy

All imaging experiments were performed on a LSM 880 Zeiss confocal microscope equipped with a water-immersion C-Apo 40X/1.2 NA. Hoechst and GFP/AF488 were excited using 405 nm and 488 nm lasers, respectively. Fluorescence was collected between 420-470 nm for Hoechst, and 500-550 nm for GFP and AF488, both on the GaAsP detector. Laser power used for imaging was adjusted to minimize photobleaching. Pixel sizes were ranging between 50 and 100 nm depending on the experiments. For live-cell experiments, cells were maintained at 37°C using a heating chamber.

To generate local 8-oxoG lesions by micro-irradiation, cell nuclei were irradiated once within a region of interest (ROI) of 80-pixel wide and 5-pixel high with a Ti:Saphirre femtosecond infrared laser (Mai Tai HP, Spectra Physics) which emission wavelength was set to 800 nm. Protein recruitment at sites of micro-irradiation was monitored by timelapse acquisitions at 1.6 Hz for approximately 90s.

For the fluorescence recovery after photobleaching (FRAP) experiments, GFP-tagged constructs were bleached within a given ROI of the nucleus using 5 iterations of the 488 nm laser set at full power and the fluorescence recovery was imaged for 10 to 25s at 13 Hz acquisition speeds using a pixel size of 208 nm. For the FRAP within the DNA damage region, the same area was bleached before and approximately 10 seconds after micro-irradiation. DNA damage was induced within a ROI of 55 by 25 pixels as described above and the FRAP ROI was either a 15-pixel wide circle or a 16-pixel wide square within the micro-irradiation ROI.

For the fluctuation correlation spectroscopy (FCS) experiments, single photons emitted by GFP-tagged proteins were counted on the GaAsP detector in photon-counting mode. Each acquisition lasted 30 seconds to minimize noise on the autocorrelation curves. They were performed before micro-irradiation and approximately 10s after damage induction.

### Image analysis

All image quantifications were performed using Fiji. For the colocalization analysis, Pearson’s correlation coefficients between the two channels were computed using the Coloc 2 Fiji plugin (https://imagej.net/plugins/coloc-2). For each cell, the Pearson’s coefficient was estimated within a manually chosen area encompassing each nucleus. To quantify protein recruitment at sites of damage, mean fluorescence signals were measured within the micro-irradiated area (*I*_*MIA*_), the whole nucleus (*I*_*Nuc*_) and in a background region (*I*_*BG*_). The intensities with the irradiated area were corrected from background and imaging photobleaching, and normalized to the mean intensity prior to laser micro-irradiation as follows :

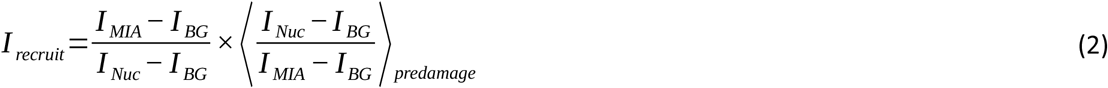

From these intensity measurements, peak recruitment values were extracted. For each acquisition, the peak intensity was considered to be the mean of all the values equal or above 95% of the absolute maximum. Peak recruitment values were estimated within the first 30 sec following damage induction. The retention of fluorescently tagged OGG1 mutants at the sites of damage was compared to that of WT OGG1 using the following metric (Fig S1A). From the recruitment curves of untreated WT OGG1, we estimated the mean time of the recruitment peak (*t*_*max*_) as well as the mean time at which half of this peak is dissipated (*t*_*1/2*_). Then, the relative residual intensity at sites of damage is estimated at *t*_*1/2*_ for the different conditions as follows:

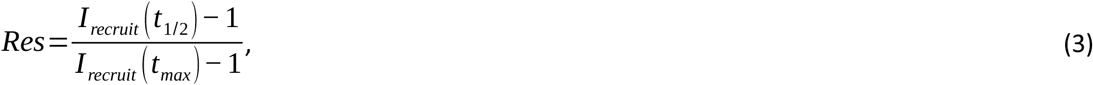

where *I*_*recruit*_(*t*_*1/2*_) and *I*_*recruit*_(*t*_*max*_) are the recruitment intensities estimated at *t*_*1/2*_ and *t*_*max*_, respectively. To analyze the FRAP acquisitions with variable sizes of photobleaching area, the intensities within the bleached region were corrected for background and imaging photobleaching similarly as what was done to quantify recruitment intensities. For the FRAP at DNA lesions, the intensities were measured in two neighboring regions both located within the micro-irradiated area: the bleached region and a unbleached reference region (Fig S1B). After background subtraction, the ratio between the intensity in the bleached and reference regions was estimated. Then, for all FRAP analysis, we subtracted the signal estimated immediately after the bleach and normalized the intensities to the pre-bleach signal. These normalized FRAP curves were fitted with the following model :

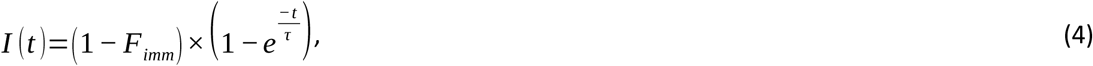

with *F*_*imm*_ the immobile fraction and τ the characteristic recovery time (35).

### Analysis of the fluorescence correlation spectroscopy data

Raw time traces were autocorrelated using the Fluctuation Analyzer 4G software after detrending for slow fluctuations using a detrending frequency of 0.125 Hz (36). When characterizing the exploration dynamics of WT OGG1 within the nucleus, the autocorrelation curves were fitted with the following diffusion model (26):

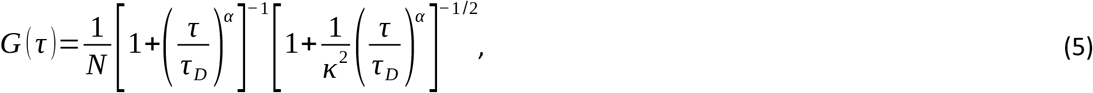

where N is the number of fluorescent protein within the confocal volume, *τ*_*D*_ is the characteristic residence time within the confocal volume, *α* is the anomalous parameter and *κ* is the structural parameter, which was fixed to 6. *α* was either fixed to 1 to fit the curves with a simple diffusion model or left free for the fitting with the anomalous diffusion model. When comparing the dynamics of different OGG1 constructs, the curves were fitted with a simple diffusion model (*α* fixed to 1) to estimate their relative residence times within the confocal volume.

### Statistics

Statistics, figures and quantification post-treatments were performed using python home-made routines. Statistics were performed using the statannot (https://pypi.org/project/statannot/) module from python. For the recruitment, FRAP and FCS curves, median±SD are shown. Unless stated otherwise, all experiments were performed in at least three independent replicates and when data from a single experiment is shown they correspond to a representative replicate. For the boxplots, the bold line indicates the median value, the box limits correspond to the 25th and 75th percentiles and the whiskers extend 1.5 times the interquartile range. p values were calculated with an independent t-test when comparing the Pearson’s coefficients, with the Wilcoxon test when comparing the acquisitions pre and post DNA damage within the same nuclei, and with the Mann-Whitney test when comparing different conditions within a given experiment. On the boxplots, * refers to p<0.05, ** to p<0.01, *** to p<0.001, **** to p<0.0001 and ns to non significant.

## RESULTS

### OGG1 alternates between transient binding to DNA and free diffusion when exploring the nuclear space in the absence of external stress

*In vitro* data have shown that OGG1 dynamically scans naked DNA during its search for 8-oxoG (21, 37). Nevertheless, it is less clear how OGG1 explores the dense environment of the nucleus in which DNA is packed within a highly folded chromatin structure. To analyze the behavior of OGG1 in living cells, we expressed wild-type (WT) OGG1 (or point mutants later in this report) fused to GFP in HeLa cells where the endogenous *OGG1* gene was knocked-out (KO) by CRISPR/Cas9 (Fig S2A,B). First, we monitored the dynamics of OGG1 within the nucleus using Fluorescence Correlation Spectroscopy (FCS), a method which allows protein motions to be characterized at high time-resolution (26). Based on the estimation of its residence times within the confocal volume, OGG1-GFP appeared much slower than two purely diffusive tracers: a GFP dimer (GFP2) and a GFP pentamer (GFP5) (Fig 1A). As these two tracers have respectively similar or larger molecular masses than OGG1-GFP, the low OGG1 mobility cannot be explained by differences in molecular sizes unless OGG1 is part of a larger complex. Nevertheless, this hypothesis seems unlikely given that, based on the Stockes-Einstein law (38, 39), this complex would need to have a molecular mass as large as 60 MDa to account for the 10-fold lower mobility observed between GFP2 and OGG1-GFP.

**Figure 1.**
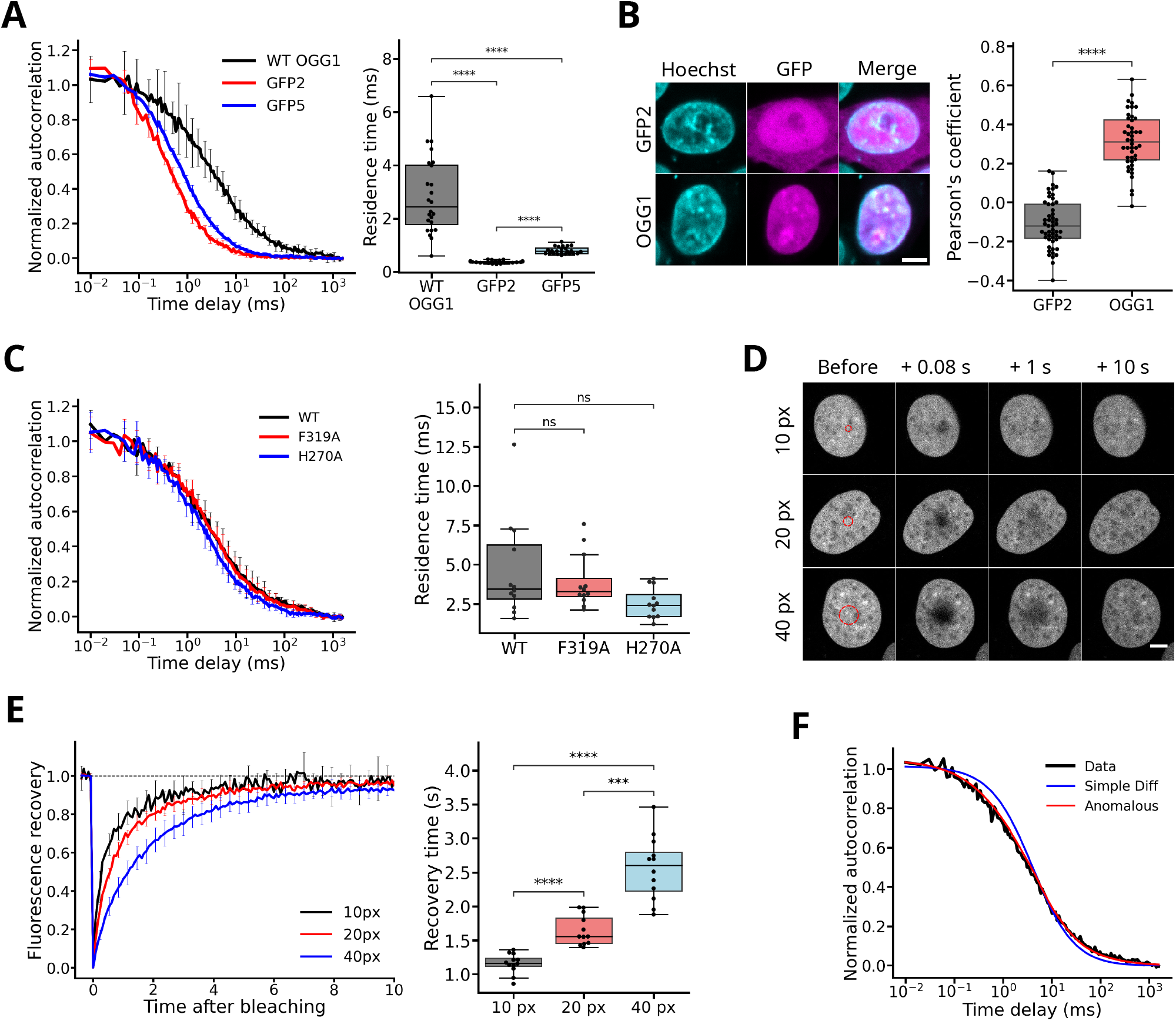
OGG1 dynamically scans the DNA within the cell nucleus. **(A)** Left: Normalized FCS autocorrelation curves obtained for WT OGG1-GFP, a GPF dimer (GFP2) and a GFP pentamer (GFP5) expressed in HeLa OGG1 KO cells. Right: Residence times of the GFP-tagged constructs within the focal volume estimated from the fit of the autocorrelation curves. 12 cells per condition. **(B)** Left: Representative confocal images of HeLa OGG1 KO cells expressing either WT OGG1-GFP or a GFP dimer (GFP2) and counterstained with Hoechst. Scale bar: 10 μm Right: Colocalization between the GFP and Hoechst channels estimated by calculation of the Pearson’s coefficients (44 cells for OGG1-GFP, 50 cells for GFP2) **(C)** Left: Normalized FCS autocorrelation curves obtained for GFP-tagged WT OGG1, OGG1-F319A and OGG1-H270A expressed in HeLa OGG1 KO cells. Right: Residence times of the GFP-tagged constructs within the focal volume estimated from the fit of the autocorrelation curves. 12 cells per condition. **(D)** Representative time-course images of the fluorescence recovery after photobleaching of circular area of variable sizes within the nucleus of HeLa OGG1 KO cells expressing WT OGG1-GFP. The bleached regions with diameters of 10, 20 and 40 pixels are shown with red dashed circles. Scale bar: 5 μm. **(E)** Left: Normalized fluorescence recovery curves for WT OGG1-GFP obtained from the images shown in D. Right: Characteristic recovery times estimated from the fit of the curves. 12 cells per condition. **(F)** Normalized autocorrelation curve obtained for WT OGG1-GFP expressed in HeLa OGG1 KO cells. Median of 12 cells. The experimental curve (black) is fitted either with a simple diffusion model (blue) or an anomalous diffusion model (red).

Putting aside size differences, the most likely interpretation of the relative slow dynamics of OGG1 compared to diffusive tracers of similar sizes is that the DNA glycosylase not only diffuses within the nucleoplasm but also transiently binds DNA, thus reducing its overall mobility. This is supported by *in vitro* data showing the dynamic association of OGG1 with oligonucleotides even in the absence of 8-oxoG (21). Consistently, we observed significant colocalization between OGG1-GFP and Hoechst-stained DNA on confocal images (Fig 1B). Because basal levels of 8-oxoG are present in the nucleus even in the absence of external stress, we asked whether the binding of OGG1 to DNA was related to the specific detection of the lesion, or rather reflected how OGG1 explores the nucleus by alternating 3D diffusion with transient association with DNA independently of its oxidation status. To address this question, we compared the nuclear dynamics of GFP-tagged WT OGG1 with that of mutated fusions harboring the F319A or H270A substitutions (Fig 1C), both of which lose the ability to bind 8-oxoG *in vitro* (40). The absence of a difference in residence time between the WT and the two mutant forms of OGG1 led us to conclude that the DNA-binding events displayed by OGG1 do not rely on specific recognition of 8-oxoG but rather correspond to the constant scanning of the DNA when searching for lesions.

Proteins that alternate between 3D diffusion in the nucleoplasm and transient association with DNA can follow three different regimes (41). In the first one, often referred to as a reaction-limited regime, diffusion between two binding sites can be considered as infinitely fast compared to the lifetime of the bound state, which is then the only parameter controlling protein dynamics. Conversely, in a diffusion-limited regime, the bound state is much shorter than diffusive transit between two binding sites. Finally, similar lifetimes between the bound and diffusive states would result in a mixed regime. To assess which regime better describes the behavior of OGG1 in the nucleus, we first performed fluorescence recovery after photobleaching (FRAP) experiments. The fact that the characteristic recovery time increased with the size of the bleached area (Fig 1D,E) implies that the dynamics of OGG1-GFP did not solely rely on the lifetime of the bound state but also on the progressive redistribution of the protein within the bleached area due to diffusion. This is incompatible with a reaction-limited regime (42). Next, if OGG1-GFP followed a diffusion-limited regime, we should be able to fit the FCS curves obtained for this protein with a simple diffusion model, similar to what can be done for a purely diffusive tracer such as GFP2 (Fig S2C) (43). However, the FCS traces acquired for OGG1-GFP could not be properly fitted with a simple diffusion model but rather required the use of an anomalous diffusion model (Fig 1F). This anomalous behavior discards the diffusion-limited regime and suggests that OGG1 proteins exploring the nuclear space rather follow a mixed regime for which the characteristic duration of the transient association with undamaged DNA to detect potential 8-oxoG lesions is within the same order of magnitude than the 3D diffusive transits between two binding events.

### The efficient recruitment of OGG1 to laser-induced 8-oxoG relies on the direct recognition of the lesion by the DNA glycosylase

After assessing its dynamics in the absence of external stress, we characterized the spatio-temporal behavior of OGG1 upon acute induction of 8-oxoG. We set up a micro-irradiation protocol based on the use of an 800-nm femtosecond pulsed laser, which generated 8-oxoG locally within the nucleus of HeLa cells, as confirmed by immunofluorescence staining (Fig 2A). The accumulation of OGG1-GFP at sites of damage started immediately after irradiation, with a peak recruitment at about 10 s post irradiation and was followed by a slower dissipation phase lasting a few minutes (Fig 2B). We wondered whether OGG1 proteins accumulating at sites of damage were stably bound to the lesions or kept exchanging with a diffusive pool. The rapid fluorescence recovery observed after photobleaching of a sub-region of the recruitment area revealed that the association of OGG1 with DNA at sites of damage is highly dynamic (Fig 2C,D). This recovery was nevertheless slower than prior to damage induction with a longer characteristic recovery time as well as a larger non-recovering fraction (Fig 2D), indicating that the glycosylase accumulation at the irradiated area was correlated with a tighter binding of OGG1 to DNA. This was also supported by probing the local mobility of OGG1-GFP before and after laser irradiation using FCS, which revealed a longer residence time of the glycosylase in the focal volume after damage induction (Fig 2E).

**Figure 2.**
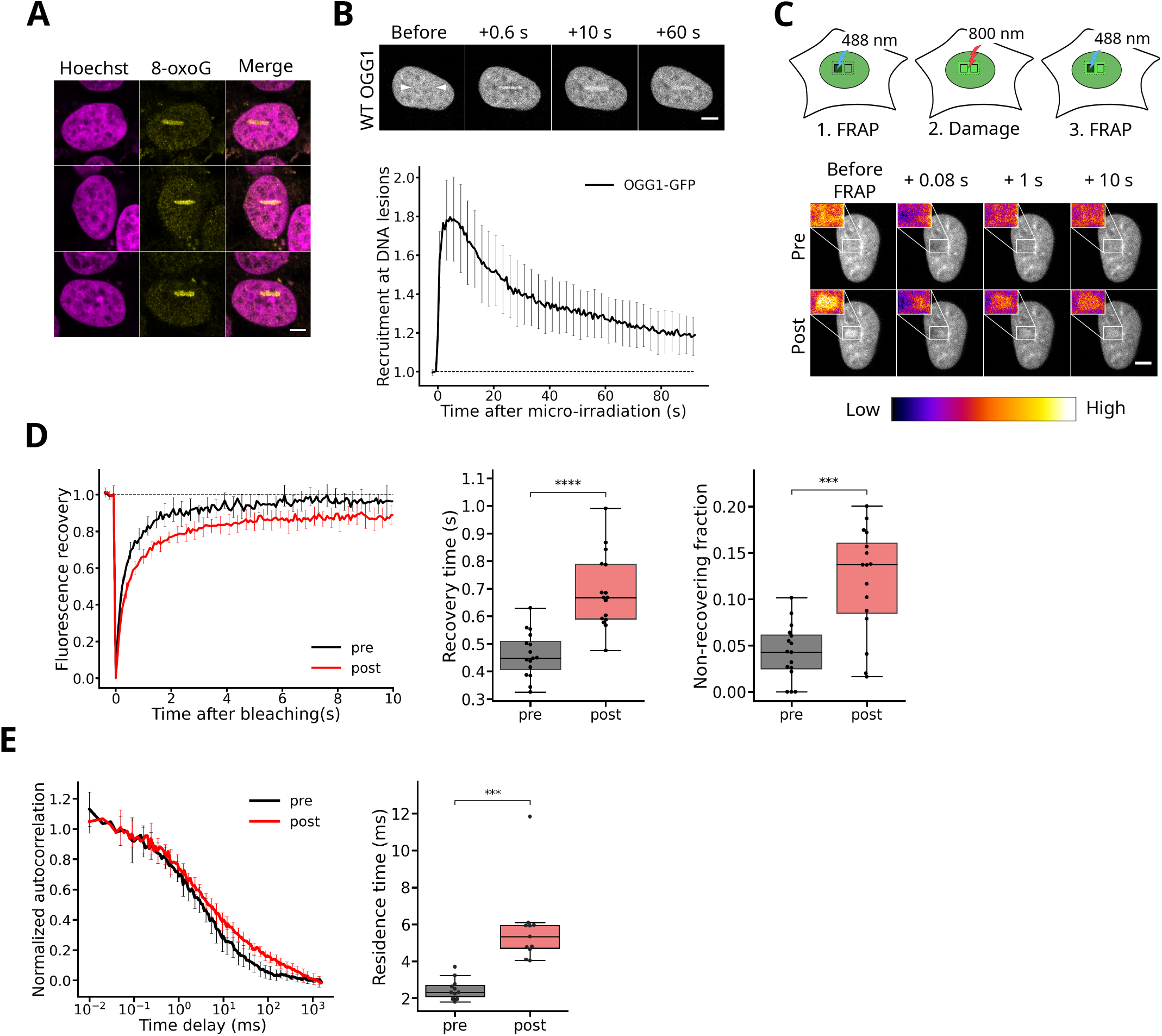
OGG1 is rapidly recruited to 8-oxoG lesions induced by laser micro-irradiation and displays rapid turnover at sites of damage. **(A)** Representative confocal images of HeLa OGG1 KO after laser micro-irradiation and immunostaining against 8-oxoG. DNA was counterstained with Hoechst. Scale bar: 5 μm **(B)** Top: Representative time-course images of the accumulation of WT OGG1-GFP at sites of laser micro-irradiation in the nucleus of OGG1 KO cells. White arrowheads indicate the micro-irradiated line. Scale bar: 5 μm. Bottom: Curve of the recruitment kinetics of WT OGG1-GFP at sites of micro-irradiation measured from the time-course images. Median of 12 cells. **(C)** Representative time-course images of the fluorescence recovery after photobleaching within the nucleus of HeLa OGG1 KO cells expressing WT OGG1-GFP before damage (pre) and at sites of laser micro-irradiation (post). Insets in pseudocolor show a magnified view of the micro-irradiated region. Scale bar: 5 μm. **(D)** Left: Normalized fluorescence recovery curves for WT OGG1-GFP before and after damage induction derived from the images shown in C. Right: Characteristic recovery times and immobile fractions estimated from the fits of the fluorescence recovery curves. 16 cells before and after damage. **(E)** Left: Normalized FCS autocorrelation curves measured for WT OGG1-GFP before damage (pre) and at sites of laser micro-irradiation (post) in the nucleus of HeLa OGG1 KO cells. Right: Residence times of WT OGG1-GFP within the focal volume estimated from the fit of the autocorrelation curves. 11 cells before and after damage.

This rapid accumulation of OGG1 at sites of damage could rely on the recognition of 8-oxoG by the glycosylase itself but also involve other unknown 8-oxoG binding factors that OGG1 would associate with. To assess the specific contribution of the direct association of OGG1 with 8-oxoG to the recruitment process, we analyzed the behavior at sites of micro-irradiation of the OGG1-F319A and H270A mutants, which both show very low affinity for 8-oxoG *in vitro* (40). Albeit not fully abrogated, the accumulation of these two mutants was significantly reduced compared to WT (Fig 3A,B), in line with the need for a direct recognition of 8-oxoG by the glycosylase for efficient recruitment. Because these mutants have lost affinity for 8-oxoG but are still able to bind to apurinic/apyrimidinic (AP) sites (40), we asked whether their residual recruitment at the sites of DNA damage could be due to their binding to AP sites generated by the laser irradiation concomitantly to 8-oxoG. We found that the endonuclease APE1, which processes AP sites, was recruited to the sites of damage as early as OGG1 (Fig S3A), suggesting that laser irradiation produced AP sites. In addition, the differences in peak recruitment between the three OGG1 mutants F319A, H270A and H270L (Fig S3A,B), which all lost ability to bind 8-oxoG, correlated with their decreasing affinity for the AP-site analog THF (40). These two findings suggest that the accumulation observed for WT OGG1 within the irradiated area is partially due to the presence of AP-sites, which could also explain the incomplete abrogation of the recruitment of the OGG1 mutants unable to bind to 8-oxoG. In addition to the analysis of protein recruitment, we also assessed protein dynamics at sites of laser irradiation by FCS. In contrast to what was observed for WT OGG1, we found no change in the local mobility for OGG1-F319A nor H270A compared to undamaged conditions (Fig 3C), consistent with their reduced affinity for the lesions generated at sites of laser micro-irradiation.

**Figure 3.**
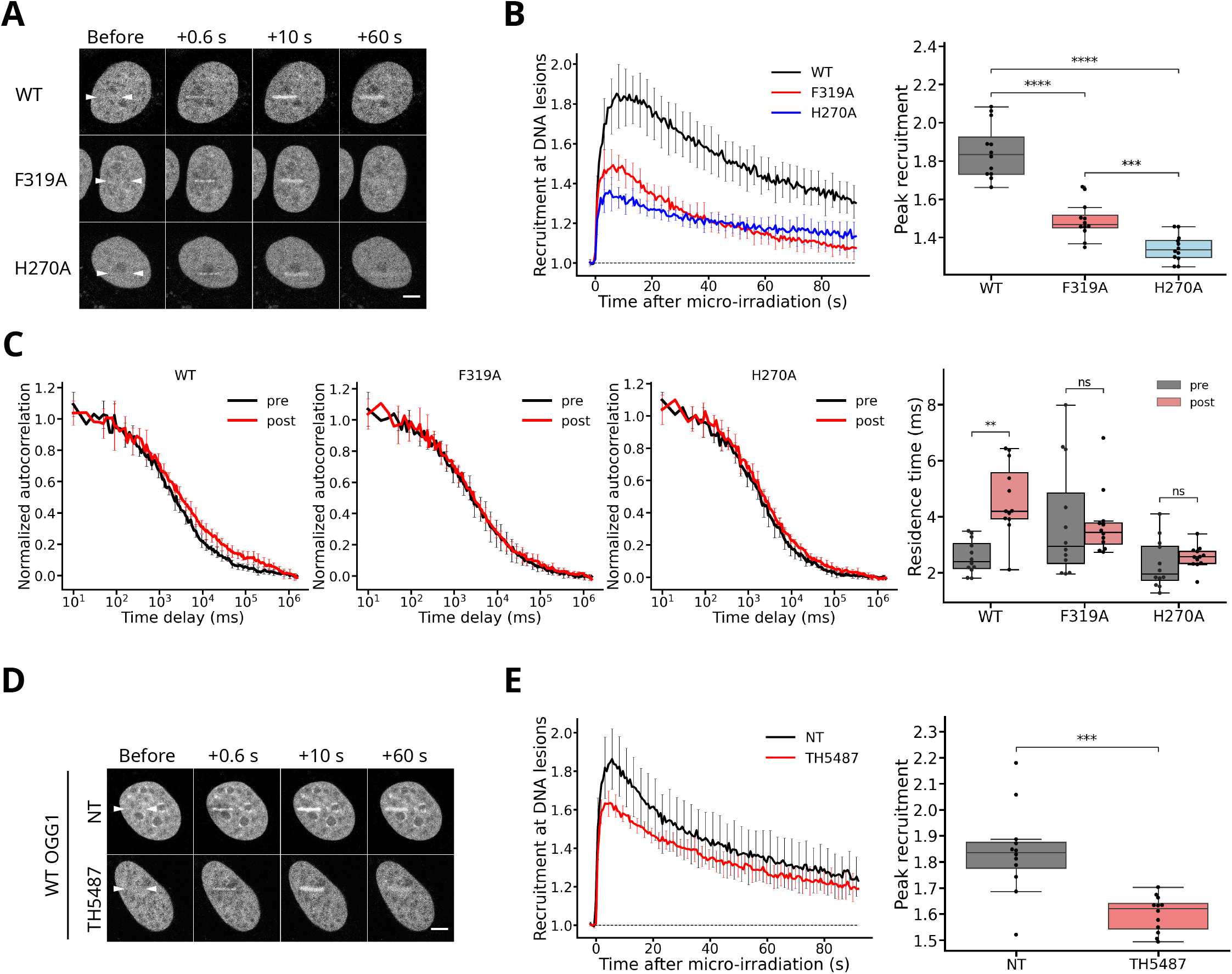
Direct detection of 8-oxoG by OGG1 is essential for its efficient accumulation at sites of DNA damage. **(A)** Representative time-course images of the accumulation of GFP-tagged OGG1-WT, OGG1-F319A and OGG1-H270A at sites of laser micro-irradiation in the nucleus of HeLa OGG1 KO cells. White arrowheads indicate the micro-irradiated line. Scale bar: 5 μm. **(B)** Left: Curves of the recruitment kinetics of OGG1-WT, F319A and H270A at sites of micro-irradiation derived from the images shown in A. Right: Peak recruitment extracted from the recruitment curves. 12 cells per condition. **(C)** Left: Normalized FCS autocorrelation curves measured for GFP-tagged OGG1-WT, OGG1-F319A and OGG1-H270A before damage (pre) and at sites of laser micro-irradiation (post) in the nucleus of HeLa OGG1 KO cells. Right: Residence time of the different constructs within the focal volume estimated from the fit of the autocorrelation curves. 12 cells per condition. **(D)** Representative time-course images of the accumulation of WT OGG1-GFP at sites of laser micro-irradiation in the nucleus of HeLa OGG1 KO cells left untreated (NT), or treated with 30 μM of the OGG1 inhibitor TH5487. White arrowheads indicate the micro-irradiated line. Scale bar: 5 μm. **(E)** Left: Curves of the recruitment kinetics of OGG1-WT at sites of micro-irradiation derived from the images shown in D. Right: Peak recruitment extracted from the recruitment curves. 12 cells per condition.

To confirm our conclusions derived from the use of OGG1 mutants, we also analyzed the impact of the OGG1 competitive inhibitor TH5487 (44, 45), on the recruitment of the glycosylase to sites of DNA damage induced by laser micro-irradiation. In line with the fact that TH5487 binds to the active site of OGG1, thus precluding association of the glycosylase with 8-oxoG, we observed a significant reduction in OGG1 accumulation to sites of damage in cells treated with this inhibitor (Fig 3D,E). These results are fully consistent with those obtained with the OGG1 mutants F319A and H270A, and demonstrate that the direct recognition of 8-oxoG by OGG1 is crucial for the rapid accumulation of the glycosylase to DNA lesions.

### The conserved residue G245 is essential for OGG1 association with DNA

In the previous sections, we described the use of different fluorescence-based methods to assess two specific aspects of OGG1 behavior in living cells: i) how the glycosylase explores the nuclear space when searching for its cognate lesion and ii) how it efficiently accumulates at DNA lesions induced by laser micro-irradiation. Next, we used these assays to analyze the potential involvement of specific residues of OGG1 on these two components of its nuclear dynamics. We first focused on the residue G245, which belongs to the helix/hairpin/helix domain and is thought to mediate association of OGG1 with DNA (17, 19). While structural data have shown that this amino-acid contacts the DNA backbone approximately 3 bp away from the lesion (17), the exact function of this highly conserved residue remains poorly defined. By EMSA, the purified OGG1-G245A mutant displayed no detectable affinity for DNA duplexes harboring or not an 8-oxoG (Fig 4A). In cells, GFP tagged OGG1-G245A showed higher mobility than the WT protein in the absence of induced damage (Fig 4B). Together, these findings highlight the critical role of the G245 residue in OGG1 interaction with DNA irrespectively of the presence of 8-oxoG, and confirm that transient association with DNA is a key factor regulating the exploration of the nucleus by OGG1. Furthermore, in line with the impaired affinity for 8-oxoG observed *in vitro*, we also found that OGG1-G245A displayed no detectable catalytic activity (Fig 4C,D) and was unable to accumulate at DNA lesions induced by laser irradiation (Fig 4E,F).

**Figure 4.**
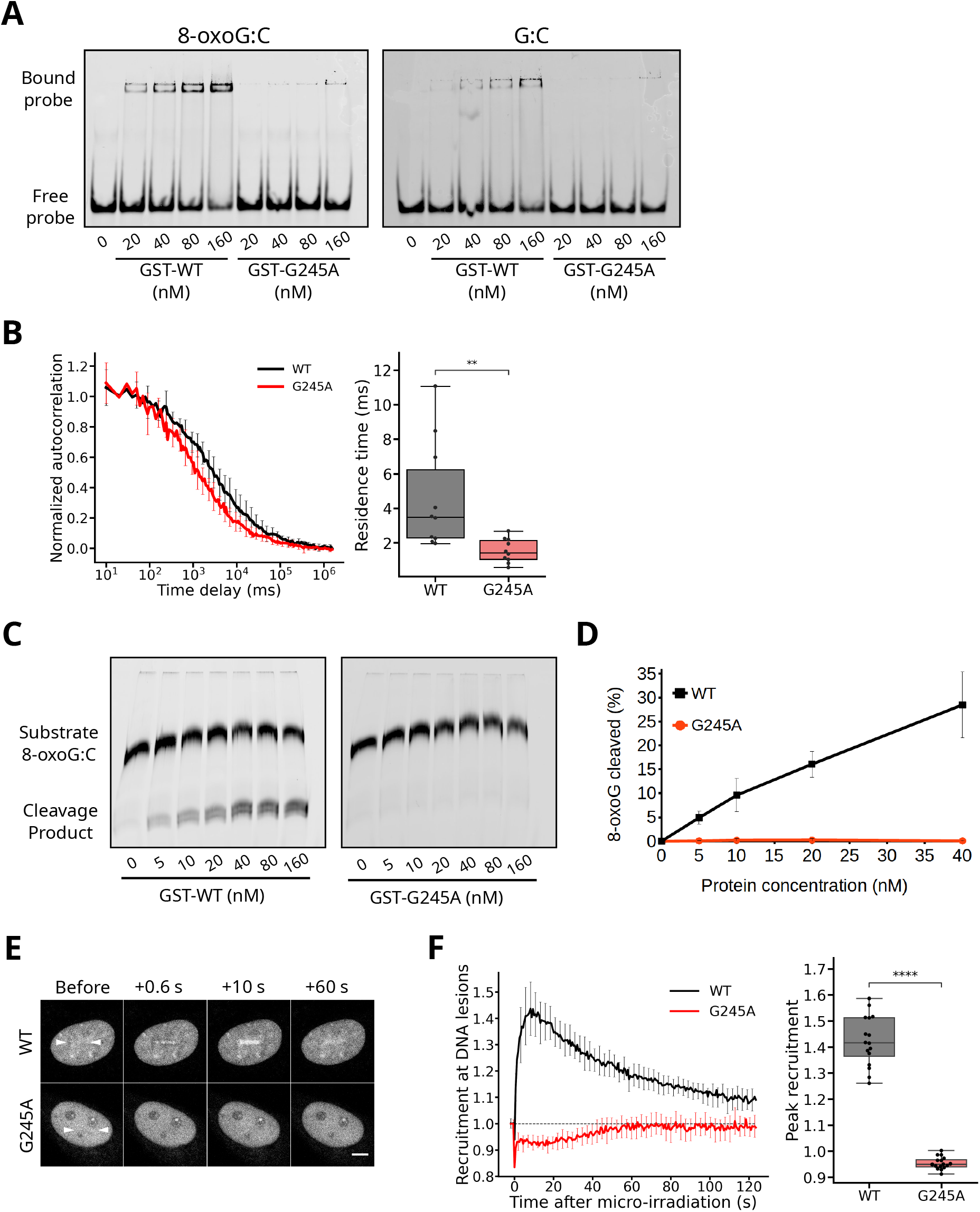
Conserved residue G245 is essential for OGG1 association with DNA. **(A)** Representative gel-shifts showing the binding of purified GST-tagged OGG1-WT and OGG1-G245A to 8-oxoG:C (left) and G:C (right) containing DNA duplexes for concentrations of proteins ranging between 0 and 160 nM. **(B)** Left: Normalized FCS autocorrelation curves measured for GFP-tagged OGG1-WT and OGG1-G245A in the absence of laser micro-irradiation in the nucleus of HeLa OGG1 KO cells. Right: Residence time of the two constructs within the focal volume estimated from the fit of the autocorrelation curves. 10 cells per condition. **(C)** Representative gels of the cleavage of an 8-oxoG:C containing oligonucleotide by increasing concentrations of GST-tagged OGG1-WT and OGG1-G245A ranging between 0 and 160 nM. **(D)** Quantification of the relative amounts of cleavage product from the gels shown in C. Mean of 3 independent repeats. **(E)** Representative time-course images of the accumulation of GFP-tagged OGG1-WT and OGG1-G245A at sites of laser micro-irradiation in the nucleus of HeLa OGG1 KO cells. White arrowheads indicate the micro-irradiated line. Scale bar: 5 μm. **(F)** Left: Curves of the recruitment kinetics of OGG1-WT and OGG1-G245A at sites of micro-irradiation derived from the images shown in E. Right: Peak recruitment extracted from the recruitment curves. 16 cells per condition.

### OGG1 residues involved in base probing differentially impact transient binding to undamaged DNA and accumulation at oxidized bases

The initial steps of the inspection of the oxidation status of the guanine by OGG1 are only partially understood. While previous reports suggest that at least a partial extrahelical extrusion of the probed base promoted by DNA kinking is needed (17), recent work proposed that the discrimination between non-oxidized Gs and 8-oxoGs could occur intrahelically (19). Here, we focused on the roles of the OGG1 residues Y203, N149 and N150 in these early steps of the lesion recognition. Structurally, these residues seem to act as a two-arm clamp, one arm made of residue Y203 and the second one by amino-acids N149 and N150. This clamp is initially closed with its tip, composed of residues Y203 and N149, inserted in the DNA to promote G:C or 8-oxoG:C pair melting required for base extrusion. Although it remains unclear whether this extrusion depends on the oxidation status of the G, the base flip out is associated with the opening of the clamp which holds in place the kinked double-helix and, in particular, the estranged C (19). To better understand the role of this clamp, we analyzed the consequences of mutating each of its two arms.

First, we assessed the behavior of the OGG1 mutants Y203A and N149A/N150A (herein referred to as 2NA) *in vitro*. Compared to WT, the Y203A mutant had similar affinity to lesion-free DNA duplexes while the 2NA mutant showed dramatically increased binding (Figs 5A,B, S4 and Table 1). In contrast, both mutants displayed reduced affinity for 8-oxoG containing DNA duplexes, in particular the Y203A mutant (Fig 5A,B, S4 and Table 1). Of note, the non-specific complex between OGG1 and DNA irrespectively of the presence of the lesion (C2 complex) seemed to be characterized by a different stoichiometry compared to the specific association between the glycosylase and 8-oxoG-containing DNA duplexes (C1 complex). This might indicate that several OGG1 molecules can bind simultaneously to the same unmodified oligonucleotide at the high concentrations of glycosylase required to form the C2 complex. Interestingly, the Y203A mutant preferentially formed the non-specific complex C2 with DNA even in the presence of 8-oxoG, showing that the Y203 residue is essential for lesion recognition. We also found that both the Y203A and 2NA mutants showed dramatically reduced cleavage activity, in line with their impaired binding to 8-oxoG (Fig 5C).

**Figure 5.**
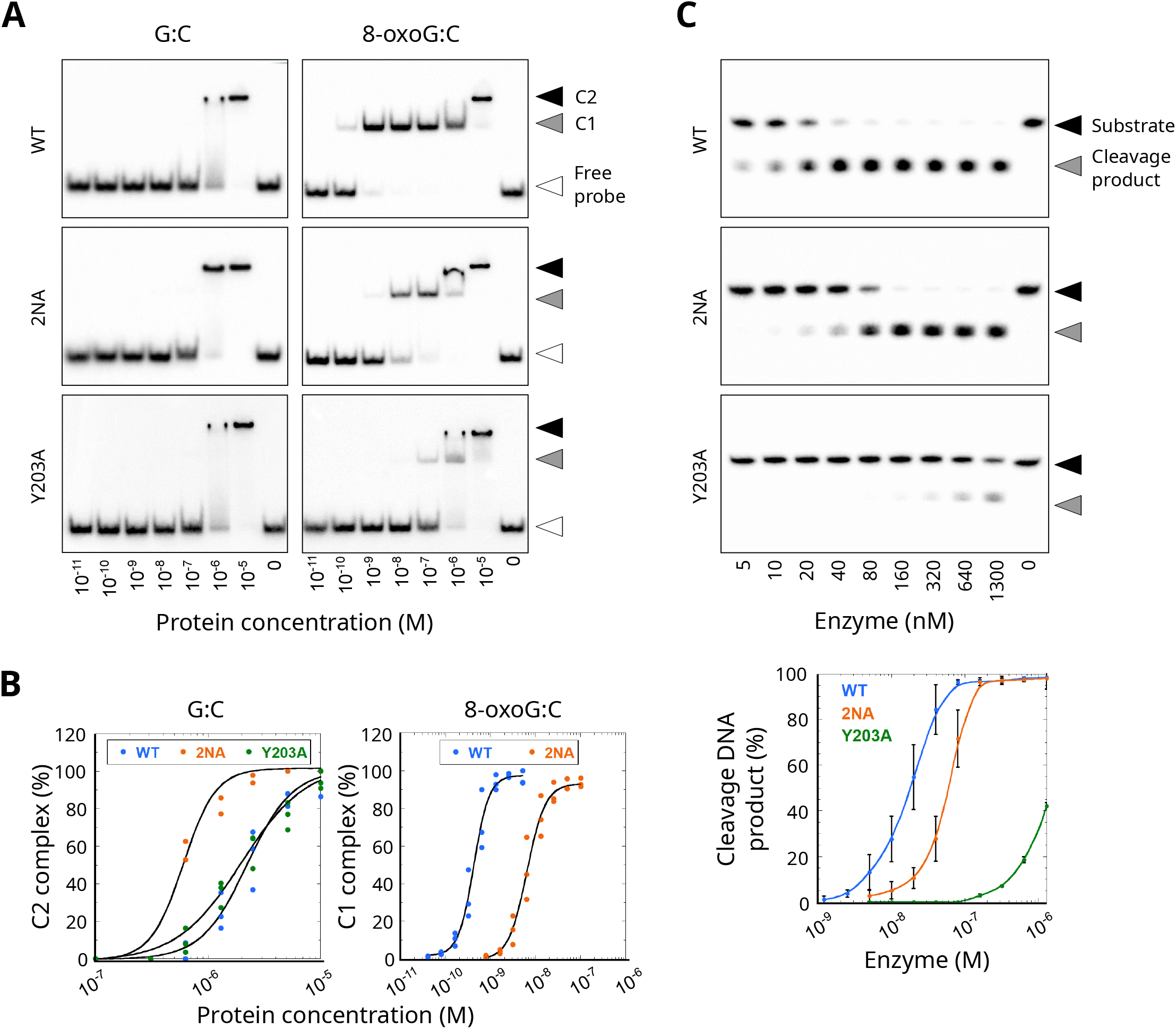
Mutating the probing residues Y203 and N149/N150 have differential impacts on the DNA binding properties and DNA-glycosylase activity of OGG1 in vitro. **(A)** Representative gel-shifts showing the binding of purified OGG1-WT, OGG1-N149A/ N150A (2NA) and OGG1-Y203A to radiolabeled DNA duplexes free of damage (G:C) or containing an 8-oxoG:C pair (8-oxoG:C). The protein concentrations are shown below the gels. C1 refers to the specific lesion recognition complex composed of 1 protein per 8-oxoG:C probe while C2 corresponds to a non-specific complex probably composed of two OGG1 proteins binding to one DNA duplex molecule, irrespectively of the presence of the lesions. **(B)** Titration curves of the relative amounts of DNA/protein complex (C1 or C2 as indicated) as a function of the protein concentration for OGG1-WT, OGG1-2NA and OGG1-Y203A interacting with an undamaged DNA probe (G:C) or an 8-oxoG:C containing DNA probe (8-oxoG:C). These titration curves were estimated from the gel-shifts shown on Fig S4. Each point corresponds to the mean of at least three independent repeats. For OGG1-Y203A, the dramatic loss of affinity for 8-oxoG did not allow to monitor the titration curve of the C1 complex. **(C)** Top: Representative gels of the amounts of 8-oxoG:C containing radiolabelled oligonucleotide substrate (S) and its OGG1 cleavage product (P) for growing concentrations of OGG1-WT, OGG1-2NA and OGG1-Y203A. Bottom: Quantification of the relative amounts of cleavage product as a function of the protein concentration from the gels shown above. Each point corresponds to the mean of at least three independent repeats.

**Table 1.** Apparent dissociation constants of purified OGG1-WT, OGG1-N149/N150A (2NA) and OGG1-Y203A bound to DNA duplexes free of damage (G:C) or containing an 8-oxoG:C base pair (8-oxoG:C). The apparent dissociation constants were estimated as the protein concentration for half maximal binding from the titration curves shown on figure 5B. Because no proper titration curve of the C1 complex could be obtained for OGG1-Y203A, we were only able to estimate a lower bound of the K_Dapp_ for this mutant based on the visual inspection of the gel-shifts on figure S4.

| OGG1 | Apparent dissociation constants (nM) |  |  |
| --- | --- | --- | --- |
|  | G:C* | 8-oxoG:C<sup>#</sup> | Ratio G:C/8-oxoG:C |
| **WT** | 2220 ± 220 | 0.42 ± 0.03 | 5286 |
| **2NA** | 602 ± 42 | 6.37 ± 0.58 | 95 |
| **Y203A** | 1970 ± 270 | > 340 | < 6 |

\*K<sub>Dapp</sub> for C2 complex (unspecific complex between OGG1 and the DNA duplexes)
<sup>#</sup>K<sub>Dapp</sub> for C1 complex (specific recognition complex between OGG1 and 8-oxoG)

Next, we analyzed the impact of the Y203A and 2NA mutations on the behavior of OGG1 in living cells. In the absence of induced damage, the nuclear mobility of the Y203A mutant was similar to that of the WT (Fig 6A), while the 2NA mutant was drastically slower (Fig 6B), indicating a stronger interaction with undamaged DNA for the latter. These results are fully consistent with the relative affinities of these two mutants for lesion-free DNA duplexes (Fig 5A,B, S4). Importantly, FRAP recovery time measured for the 2NA mutant still increased with the size of the bleached area (Fig S5A,B) suggesting that mutating residues N149/N150 did not lead to a switch of the OGG1 nuclear exploration regime. Therefore, similar to wild-type, the 2NA mutant keeps rapidly alternating between a DNA-bound and a diffusive state but mutating these two residues tends to displace the equilibrium towards the bound state. Upon laser micro-irradiation, we observed only minor accumulation of OGG1-Y203A compared to WT (Fig 6C,D), in line with its loss of ability to recognize 8-oxoG *in vitro* (Fig 5A,B, S4). The 2NA mutant was still able to accumulate in micro-irradiated regions although at lower level than WT OGG1 (Fig 6C,D), which is also consistent with the reduced binding of this mutant to the 8-oxoG containing DNA duplexes (Fig 5A,B, S4). Interestingly, OGG1-2NA displayed persistent accumulation in the damaged area, a feature that was also observed for OGG1-K249Q (Fig 6C-E), a well characterized mutant of OGG1 which binds to 8-oxoG but is unable to excise it (25, 40, 46). Despite this persistent accumulation, both mutants still displayed rapid turnover at sites of damage, similar to what was observed for WT OGG1 (Fig S5C). We also found that the release of WT OGG1 was impaired in cells treated with O8-Cl, a small molecule inhibitor that blocks OGG1 catalytic activity without affecting its binding to 8-oxoG (Fig 6F,G) (47, 48). Given that the glycosylase activity of OGG1-2NA was severely impaired (Fig 5C), these different results consistently show that the release of OGG1 from the damaged area is correlated with the progressive clearance of the 8-oxoG. This agrees with what has been shown after cell treatment with oxidizing agents (46).

**Figure 6.**
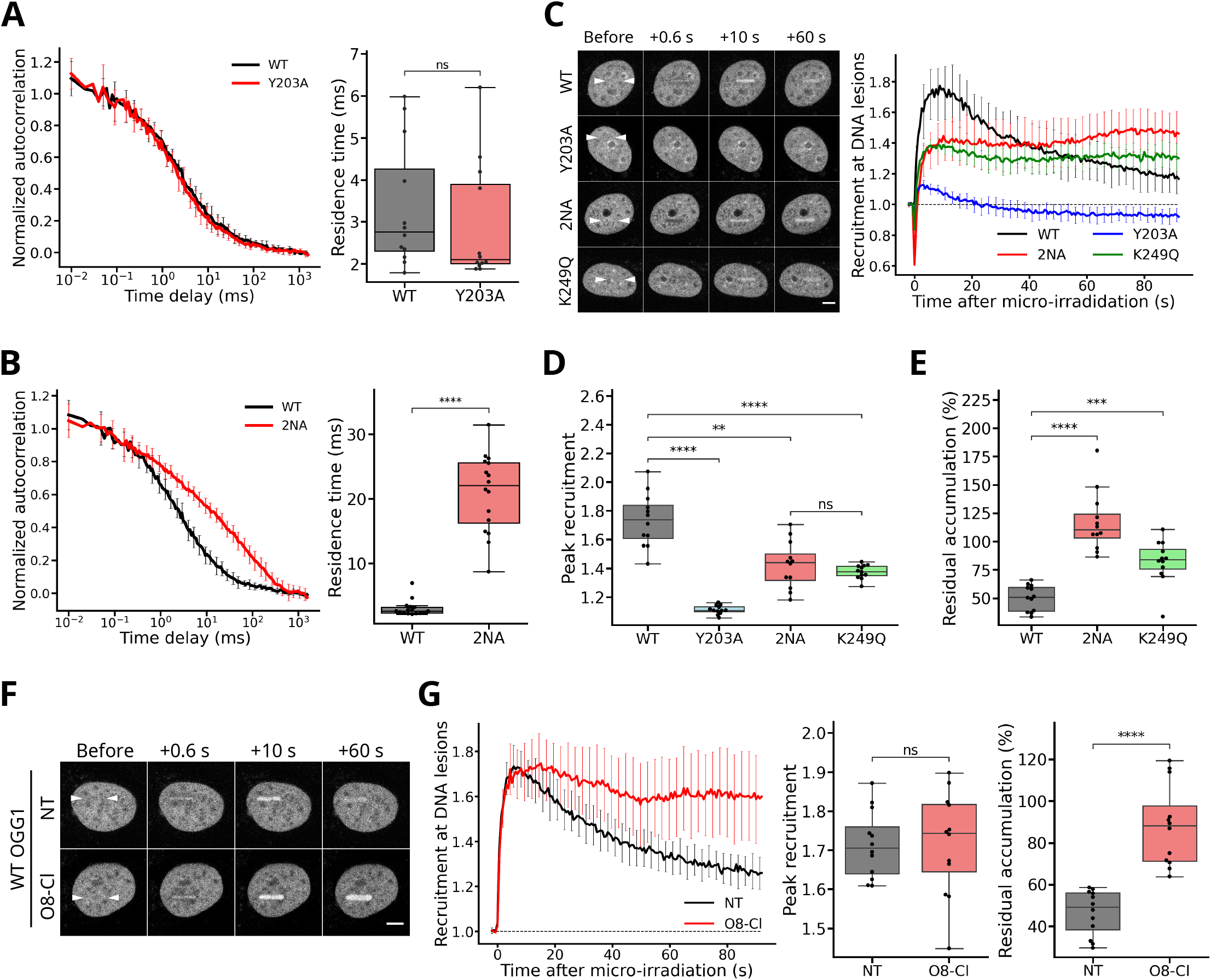
Mutating the probing residues Y203 and N149/N150 have differential impacts the dynamics of OGG1 DNA scanning and recruitment to laser-induced 8-oxoG in living cells. **(A)** Left: Normalized FCS autocorrelation curves for GFP-tagged OGG1-WT and OGG1-Y203A in the absence of laser micro-irradiation in the nucleus of HeLa OGG1 KO cells. Right: Residence time of the two constructs within the focal volume estimated from the fit of the autocorrelation curves. 12 cells per condition. **(B)** Left: Normalized FCS autocorrelation curves for GFP-tagged OGG1-WT and OGG1-N149A/N150A (2NA) in the absence of laser micro-irradiation in the nucleus of HeLa OGG1 KO cells. Right: Residence time of the two constructs within the focal volume estimated from the fit of the autocorrelation curves. 16 cells per condition. **(C)** Left: Representative time-course images of the accumulation of GFP-tagged OGG1-WT, OGG1-Y203A, OGG1-2NA and OGG1-K249Q at sites of laser micro-irradiation in the nucleus of HeLa OGG1 KO cells. White arrowheads indicate the micro-irradiated line. Scale bar: 5 μm. Right: Curves of the recruitment kinetics of the different OGG1 constructs at sites of micro-irradiation derived from the images shown on the left. **(D)** Peak recruitment of the different OGG1 constructs extracted from the recruitment curves shown in C. 12 cells per condition. **(E)** Residual accumulation relative to peak recruitment estimated for OGG1-WT, OGG1-2NA and OGG1-K249Q from the recruitment curves shown in C. This relative residual accumulation is measured at the time corresponding to the dissipation of half of the peak recruitment for the wild-type construct. 12 cells per condition. **(F)** Representative time-course images of the accumulation of WT OGG1-GFP at sites of laser micro-irradiation in the nucleus of HeLa OGG1 KO cells left untreated (NT), or treated with 30 μM of the OGG1 inhibitor O8-Cl. White arrowheads indicate the micro-irradiated line. Scale bar: 5 μm. **(G)** Left: Curves of the recruitment kinetics of OGG1-WT at sites of micro-irradiation derived from the images shown in F. Right: Peak recruitment and residual accumulation extracted from the recruitment curves. 12 cells per condition.

Taken together these results complement the current structural data by showing that the mutation of residues Y203 and N149/N150, both involved at early stages of the initiation and stabilization of base extrusion, has drastically different impacts on the dynamic behavior of the glycosylase inside the nucleus.

## DISCUSSION

OGG1, similarly to all other nuclear proteins looking for rare targets along the genome, faces a paradoxical challenge: it must be both rapid and selective in detecting its cognate lesion. This is particularly difficult for OGG1 since 8-oxoG is deeply buried within the double-helix, therefore requiring careful probing of each base (17). Our results by FCS and FRAP demonstrate that OGG1 is not freely diffusive within the nucleus but dynamically associates with the DNA (Fig 1). We found that the characteristic duration of each state, DNA-bound or freely diffusive, are similar, demonstrating that OGG1 explores the nuclear environment by rapidly alternating between short-lived association with DNA and periods of 3D free diffusion, in agreement with previous *in vitro* data (21, 37). Importantly, our findings also showed that the transient binding to DNA during nuclear exploration was not controlled by Y203, H270 or F319 (Figs 3C, 6A, 7), which are important for 8-oxoG detection at different steps of the OGG1/DNA association (40, 49), but rather rely on the conserved but poorly characterized G245 residue. This dynamic association with DNA mediated by G245 is also essential for OGG1 recruitment to DNA lesions and therefore, for 8-oxoG cleavage (Fig 4). Because this residue is located away from the extruded base (17), it seems unlikely that it directly contributes to the base flip-out or 8-oxoG recognition. Instead, it probably allows an initial contact to be established between OGG1 and the double-helix, therefore initiating the search along the DNA (Fig 7). The spatial resolution of our live cell assays was not sufficient to assess 1D diffusion of OGG1 along the DNA but it would be important in the future to establish whether this “landing residue” G245 could also regulate translocation reported for OGG1 along DNA oligonucleotides in single molecule analysis (21).

**Figure 7.**
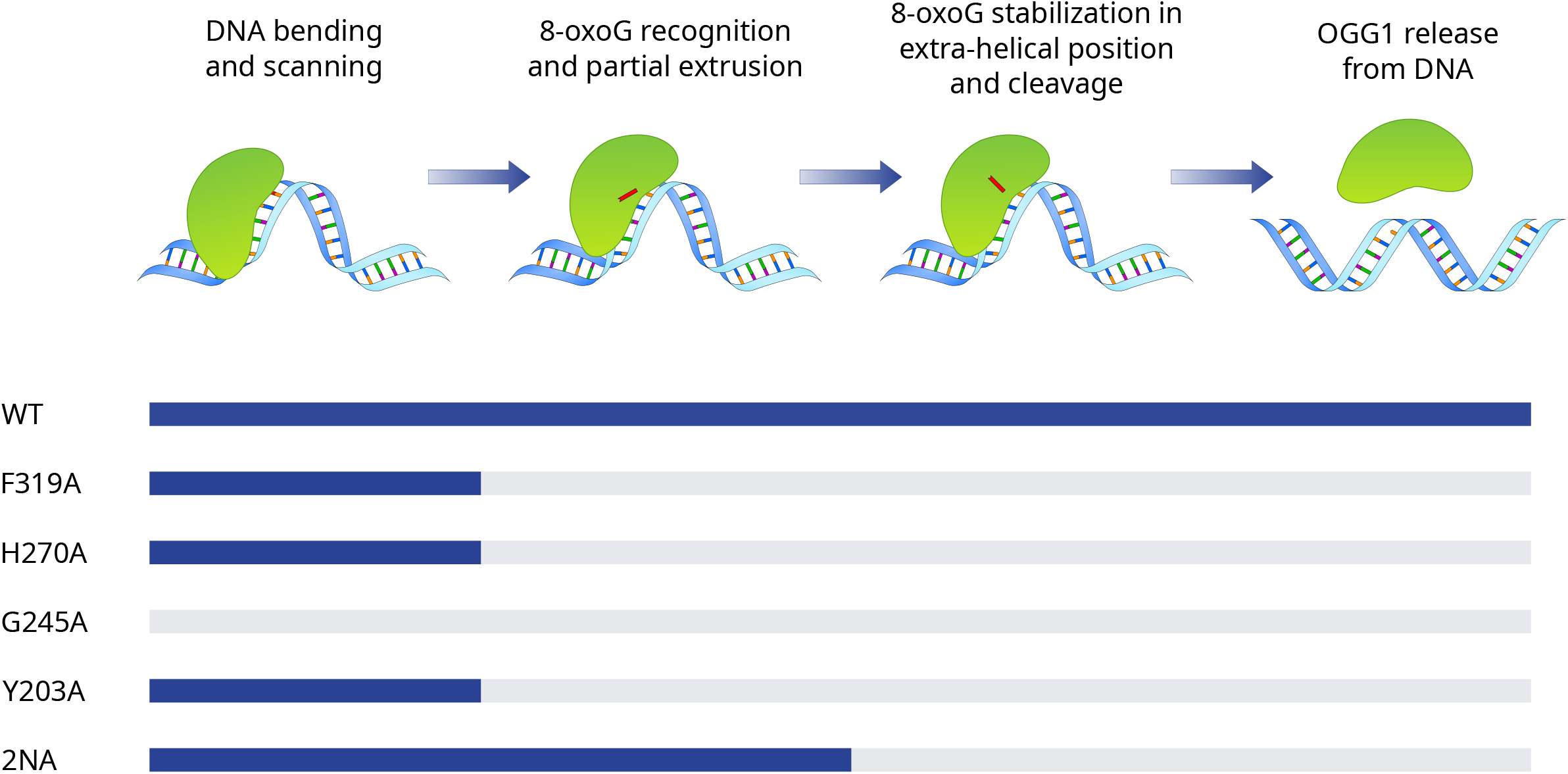
Model summarizing the impacts of the mutations studied in this work on the different steps of 8-oxoG detection and clearance by OGG1.

In agreement with previous results (17, 49, 50), we found that the residue Y203 plays a central role in the recognition of 8-oxoG, and therefore, is essential for efficient OGG1 recruitment to sites of damage (Figs 5A,B, 6A,C-D, S4). Nevertheless, structural data indicate that Y203 does not directly participate to the detection of the lesion, but rather promotes base pair destabilization (17, 51), which is probably a prerequisite for this detection (Fig 7). Then, the fact that the Y203A mutant transiently associates with undamaged DNA similarly to WT OGG1 suggests that the fraction of time spent by OGG1 for base discrimination during its transits on the DNA is small. This would be compatible with an intrahelical detection of the lesion avoiding the need for a potentially lengthy base flip-out process, as proposed recently (19). Alternatively, if extrusion is needed for base discrimination, this step should be fast compared to other processes such as OGG1 accommodation onto the DNA to promote local bending of the double helix.

The N149 and N150 residues could also contribute to base pair destabilization and flip-out by invading the void created by the extruded base and holding the distorted DNA backbone in place (17). However, provided that this base destabilization/extrusion is needed for lesion detection, our results show that residues N149 and N150 are dispensable for this process since OGG1-2NA could still recognize 8-oxoG *in vitro* and recruit to sites of damage in living cells (Figs 5A,B, 6C,D). Rather than facilitating base destabilization/extrusion, the dramatic increase in binding to undamaged DNA observed for the OGG1-2NA mutant (Figs 5A, 6B) suggests that residues N149 and N150 may help to stabilize the local structure of the DNA to avoid an over-distortion of the double-helix that could lead to a trapping of OGG1 on the DNA. As such, residues N149 and N150 would be essential to ensure the rapid reversibility of the destabilization/flip-out process in case of a normal guanine, thus allowing for a fast scanning of the DNA.

Besides these early aspects of base inspection, our results also demonstrate that OGG1 relies on its ability to directly recognize 8-oxoG to quickly accumulate at sites of DNA damage (Figs 3, S3). Furthermore, we observed a rapid turnover of the glycosylase at sites of DNA lesions, with exchange rates of few seconds based on our FRAP data (Fig 2C,D). This high turnover is also observed for OGG1 mutants that are unable to efficiently excise 8-oxoG (Fig S5C). Altogether, these data suggest that, rather than following a full sequential process starting from 8-oxoG detection, cleavage and release from the DNA, OGG1 proteins recruited to sites of damage often dissociate from the lesions before actual 8-oxoG cleavage, leading to abortive transit on the damaged DNA. Additional work will be needed to understand the mechanisms regulating this dynamic accumulation of OGG1 at DNA lesions, which probably also involve additional cofactors such as the chromatin scaffolding complex cohesin (52) or the damage sensor UV-DDB (53, 54). The approaches developed in the current work will help to further characterize the involvement of these cofactors regulating the efficiency of 8-oxoG clearance by OGG1 in the living cell context.

## DATA AVAILABILITY

The raw datasets used in the current study are available upon reasonable request.

## SUPPLEMENTARY DATA

Supplementary Data are available at NAR online.

## ACKNOWLEDGMENTS

We thank the Microscopy-Rennes Imaging Center (BIOSIT, Université Rennes 1), member of the national infrastructure France-BioImaging supported by the French National Research Agency (ANR-10-INBS-04), for providing access to the imaging setups, as well as S. Dutertre and X. Pinson for technical assistance on the microscopes. We would like to thank J. Ellenberg for sharing the plasmids coding for the GFP dimer and GFP pentamer and F. Zhang for the one coding for the GFP-tagged Cas9.

## AUTHOR CONTRIBUTIONS

O.D.A, V.G., C.S. A.M.D.G and J.P.R. completed the experiments within the manuscript. R.S., C.C. J.D., X.V., and D.B. generated cell lines and DNA constructs. J.P.R. and B.C. provided expert advice and supervised the *in vitro* experiments. A.C. and S.H. conceived and supervised this study. O.D.A, J.P.R., A.C and S.H. wrote the manuscript. All authors read and commented on the manuscript.

