## Supplemental data for "Identification of key residues of the DNA glycosylase OGG1 controlling efficient DNA scanning and recruitment to oxidized bases in living cells"

### SUPPLEMENTARY TABLES

**Table S1. List of plasmids used in this study**

| Plasmid | Source |
|---|---|
| OGG1-L1-GFP | This study |
| OGG1(F319A)-L1-GFP | This study |
| OGG1(H270A)-L1-GFP | This study |
| OGG1(H270L)-L1-GFP | This study |

| | |
|---|---|
| OGG1(G245A)-L1-GFP | This study |
| OGG1(2NA)-L1-GFP | This study |
| OGG1(Y203A)-L1-GFP | This study |
| OGG1(K249Q)-L1-GFP | This study |
| APE1-GFP | Campalans *et al.*, 2013 |
| GFP2 | Gift from Jan Ellenberg (Euroscarf P30623), Bancaud *et al.*, 2009 |
| GFP5 | Gift from Jan Ellenberg (Euroscarf P30624), Bancaud *et al.*, 2009 |
| GST-OGG1 | This study |
| GST-OGG1(G245A) | This study |
| His-OGG1 | Le Meur et al., 2015 |
| His-OGG1(Y203A) | This study |
| His-OGG1(2NA) | This study |

**Table S2. List of oligonucleotides used in this study.** Oligonucleotides are used as indicated in Material and Methods.

| **Oligonucleotide** | **Sequence (5' -> 3')** |
|---|---|
| OGG1-GFP Linker addition (For) | TCCGGAGCGGCCGCTGCAGGAGGCAGCCAAAAAATGGTGAG-CAAGGGCG |
| OGG1-GFP Linker addition (Rev) | GCCGCTCCGGATGGATCCGGGCCTTC |
| Mutation K249Q (For). In OGG1-GFP | CAGGTGGCTGACTGCATCTG |
| Mutation K249Q (Rev). In OGG1-GFP | GCAGTCAGCCACCTGGGTACCCACTCCAGGCAGGATG |
| Mutation F319A (For). In OGG1-GFP | GCCAGTGCCGACCTGCG |
| Mutation F319A (Rev). In OGG1-GFP | GGTCGGCACTGGCCAGCACCGCTTGGGC |
| Mutation G245A (For). In OGG1-GFP | CAGTGGGCACCAAGGTGG |
| Mutation G245A (Rev). In OGG1-GFP | CCTTGGTGCCCACTGCAGGCAGGATGCAGAGGG |
| Mutation H270A (For). In OGG1-GFP | GCGATGTGGCACATTGCCCAAC |
| Mutation H270A (Rev). In OGG1-GFP | CAATGTGCCACATCGCGACATCCACGGGCACAG |
| Mutation H270L (For). In OGG1-GFP | TTATGTGGCACATTGCCCAAC |
| Mutation H270L (Rev). In OGG1-GFP | GGCAATGTGCCACATAAGGACATCCACGGGCAC |
| Mutation N149A/N150A (2NA) (For). In OGG1-GFP and HisTag-OGG1 | GCGGCCAACATCGCCCGCATCAC |
| Mutation N149A/N150A (2NA) (Rev). In OGG1-GFP and HisTag-OGG1 | CGATGTTGGCCGCGGAGGAACAGATAAAAGAGAAAAGGC |
| Mutation Y203A (For). In OGG1-GFP | GTAACGGGCACGAGCGCCCAGGCCCAGC |
| Mutation Y203A (Rev). In OGG1-GFP | CTGGGCCTGGGCGCTCGTGCCCGTTAC |
| Mutation Y203A (For). In HisTag-OGG1 | GCCCGTGCCCGTTACGTG |
| Mutation Y203A (Rev). In HisTag-OGG1 | CGGGCACGGGCGCCCAGGCCCAGC |

| | |
|---|---|
| Generation of GST-OGG1 plasmids (For) | GAAAACCTTTACTTCCAGGGCCACGTGATGCCTGCCCGCG |
| Generation of GST-OGG1 plasmids (Rev) | GGGCTAGCTCTAGACTATTAGGATCCT-CAGCCTTCCGGCCCTTTG |
| Cy5-8-oxoG.DNA glycosylase activity (Fig 4) | (Cy5)-GGCTTCATCGTTGTC(8oxoG)CAGACCTGGTGGATACCG |
| Cy5-Undammaged. DNA glycosylase activity (Fig 4) | (Cy5)-GGCTTCATCGTTGTCGCAGACCTGGTGGATACCG |
| Complementary G(C)G for Cy5. DNA glycosylase activity (Fig 4) | CGGTATCCACCAGGTCTGCGACAACGATGAAGCC |
| 5'-[32P]-labeled-oxoG. DNA glycosylase activity (Fig 5) | CTGATCGATGAC(8oxoG)CCTGACATGAT |
| 5'-[32P]-labeled-undammaged. DNA glycosylase activity (Fig 5) | CTGATCGATGACGCCTGACATGAT |
| Complementary G(C)G for 5'-[32P]-labeled. DNA glycosylase activity (Fig 5) | ATCATGTCAGGCGTCATCGATCAG |
| guideRNA OGG1 Exon3 (antisense) | CACCGAAAGAGAAAAGGCATTCGAT |
| guideRNA OGG1 Exon3 (sense) | AAACATCGAATGCCTTTTCTCTTTC |

### SUPPLEMENTARY FIGURES LEGENDS

**Figure S1. Analysis of protein recruitment kinetics and fluorescence recovery after photobleaching at sites of laser micro-irradiation.** (A) To assess protein retention of OGG1 mutants at DNA lesions compared to WT, the times of the peak recruitment ($t_{max}$) and dissipation of half of the peak signal ($t_{1/2}$) were first estimated from the recruitment curves of OGG1 WT. Then, the recruitment intensity at time $t_{1/2}$ is computed as the percentage of the fluorescence intensity at $t_{max}$ for the different OGG1 constructs. (B) To estimate the fluorescence recovery at DNA lesions, the fluorescence intensities were measured in two neighboring regions within the micro-irradiated area: the photobleached region and an unbleached reference region. After background subtraction, the ratio between the intensities in the bleached and unbleached regions was estimated.

**Figure S2. Validation of the HeLa OGG1 KO cell line and analysis of the nuclear dynamics of the GFP dimer.** (A) Left: Western blot of Hela WT and OGG1 knockout cells. Vinculin is used as a loading control. Right: Quantification of the relative amount of OGG1 staining from the gel shown on the left. (B) Top: Representative gels of the amounts of 8-oxoG:C containing oligonucleotide substrate and its OGG1 cleavage product in the absence of cell extract (-) or incubated for the indicated times with extracts of WT or OGG1 KO Hela cells. Bottom: Quantification of the relative amounts of cleavage product from the gels shown above. Mean of 2 independent experiments. (C) Normalized FCS autocorrelation curve obtained in the nucleus of HeLa OGG1 KO cells expressing the GFP dimer. Median of 12 cells. The experimental curve (black) is fitted with a simple diffusion model (red).

**Figure S3. Recruitment kinetics of OGG1 mutants and APE-1 at sites of laser micro-irradiation.** (A) Curves of the recruitment kinetics for GFP-tagged OGG1-WT, F319A, H270A, H270L and AP-endonuclease APE1 expressed in HeLa OGG1 KO cells. 12 cells per condition. (B) Peak recruitment extracted from the curves shown in A for OGG1-WT, F319A, H270A and H270L.

**Figure S4. DNA binding properties of purified OGG1-WT, OGG1-N149A/N150A (2NA) and OGG1-Y203A.** Representative gel-shifts showing the binding of purified OGG1-WT, OGG1-N149A/N150A (2NA) and OGG1-Y203A to radiolabeled DNA duplexes free of lesion (G:C) or containing an 8-oxoG:C pair (8-oxoG:C). The protein concentrations are shown below the gels. The protein/DNA complexes C1 and C2 are defined in the caption of figure 5.

**Figure S5. Nuclear dynamics of the OGG1-N149A/N150A (2NA) mutant in the absence of external damage and at sites of laser micro-irradiation.** (A) Representative time-course images of the fluorescence recovery after photobleaching of circular area of variable sizes within the nucleus of HeLa OGG1 KO cells expressing GFP-tagged OGG1-WT or OGG1-2NA. The bleached regions of 15 and 30 pixel diameters are shown with red dashed circles. Scale bar: 5 µm. (B) Left: Normalized fluorescence recovery curves for GFP-tagged OGG1-WT and OGG1-2NA obtained from the images shown in A. Right: Characteristic recovery times estimated from the fit of the curves. 12 cells per condition. (C) Representative time-course images of the fluorescence recovery after photobleaching a sub-region of the area subjected to laser micro-irradiation within the nucleus of HeLa OGG1 KO cells expressing GFP-tagged OGG1-WT, OGG1-2NA or OGG1-K249Q. Insets in pseudocolor show a magnified view of the micro-irradiated region. Scale bar: 5 µm.

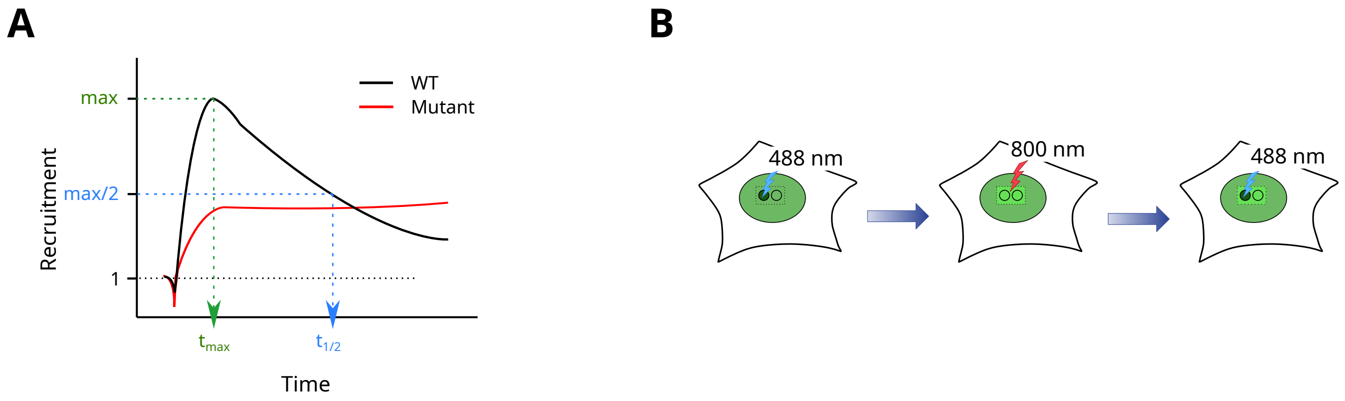


**Figure S1. Analysis of protein recruitment kinetics and fluorescence recovery after photobleaching at sites of laser micro-irradiation.** **(A)** To assess protein retention of OGG1 mutants at DNA lesions compared to WT, the times of the peak recruitment (tmax) and dissipation of half of the peak signal (t1/2) were first estimated from the recruitment curves of OGG1 WT. Then, the recruitment intensity at time t1/2 is computed as the percentage of the fluorescence intensity at tmax for the different OGG1 constructs. **(B)** To estimate the fluorescence recovery at DNA lesions, the fluorescence intensities were measured in two neighboring regions within the micro-irradiated area: the photobleached region and an unbleached reference region. After background subtraction, the ratio between the intensities in the bleached and unbleached regions was estimated.

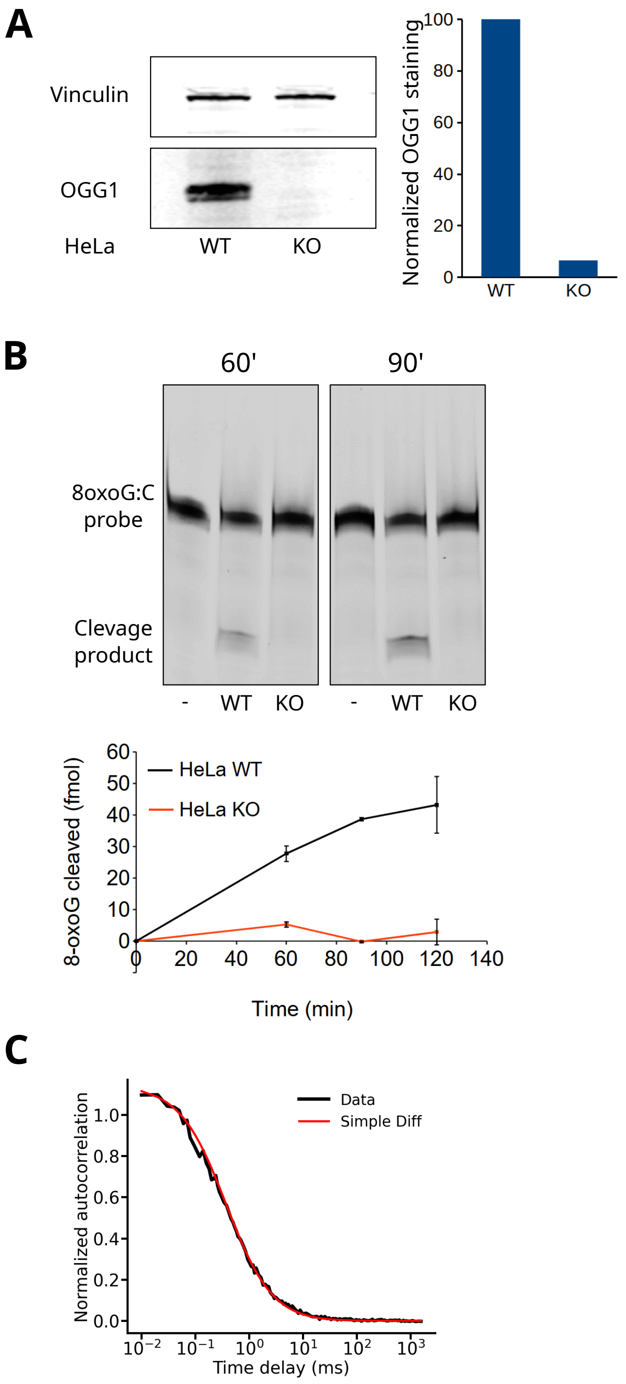


**Figure S2. Validation of the HeLa OGG1 KO cell line and analysis of the nuclear dynamics of the GFP dimer. (A)** Left: Western blot of Hela WT and OGG1 knockout cells. Vinculin is used as a loading control. Right: Quantification of the relative amount of OGG1 staining from the gel shown on the left. **(B)** Top: Representative gels of the amounts of 8-oxoG:C containing oligonucleotide substrate and its OGG1 cleavage product in the absence of cell extract (-) or incubated for the indicated times with extracts of WT or OGG1 KO Hela cells. Bottom: Quantification of the relative amounts of cleavage product from the gels shown above. Mean of 2 independent experiments. **(C)** Normalized FCS autocorrelation curve obtained in the nucleus of HeLa OGG1 KO cells expressing the GFP dimer. Median of 12 cells. The experimental curve (black) is fitted with a simple diffusion model (red).

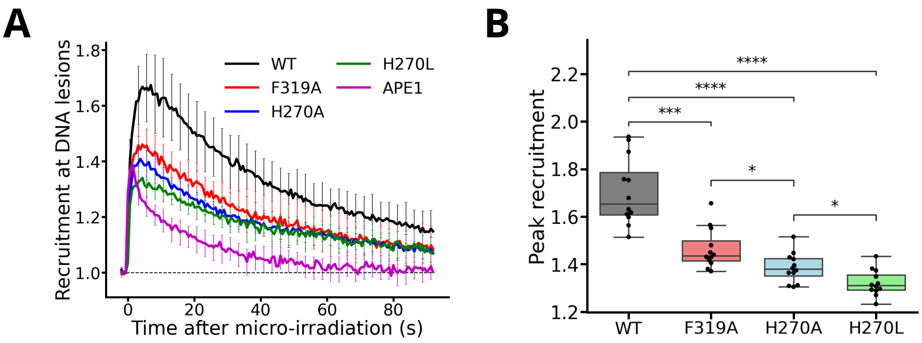


**Figure S3. Recruitment kinetics of OGG1 mutants and APE-1 at sites of laser micro-irradiation. (A)** Curves of the recruitment kinetics for GFP-tagged OGG1-WT, F319A, H270A, H270L and AP-endonuclease APE1 expressed in HeLa OGG1 KO cells. 12 cells per condition. **(B)** Peak recruitment extracted from the curves shown in A for OGG1-WT, F319A, H270A and H270L.

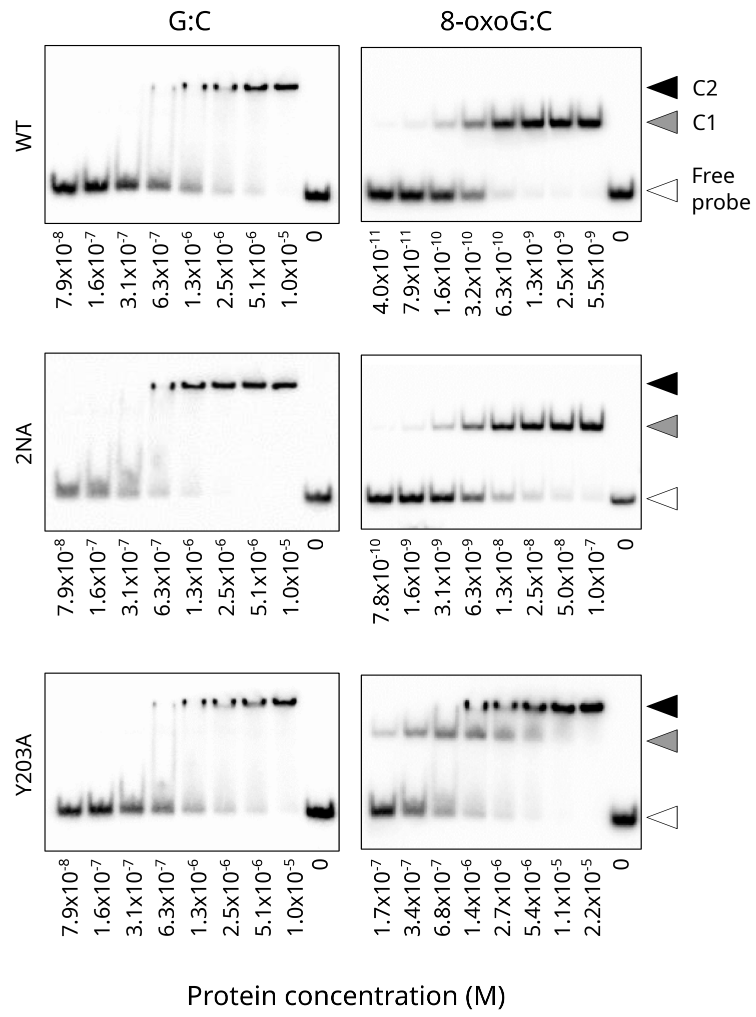


**Figure S4. DNA binding properties of purified OGG1-WT, OGG1-N149A/N150A (2NA) and OGG1-Y203A.** Representative gel-shifts showing the binding of purified OGG1-WT, OGG1-N149A/N150A (2NA) and OGG1-Y203A to radiolabeled DNA duplexes free of lesion (G:C) or containing an 8-oxoG:C pair (8-oxoG:C). The protein concentrations are shown below the gels. The protein/DNA complexes C1 and C2 are defined in the caption of figure 5.

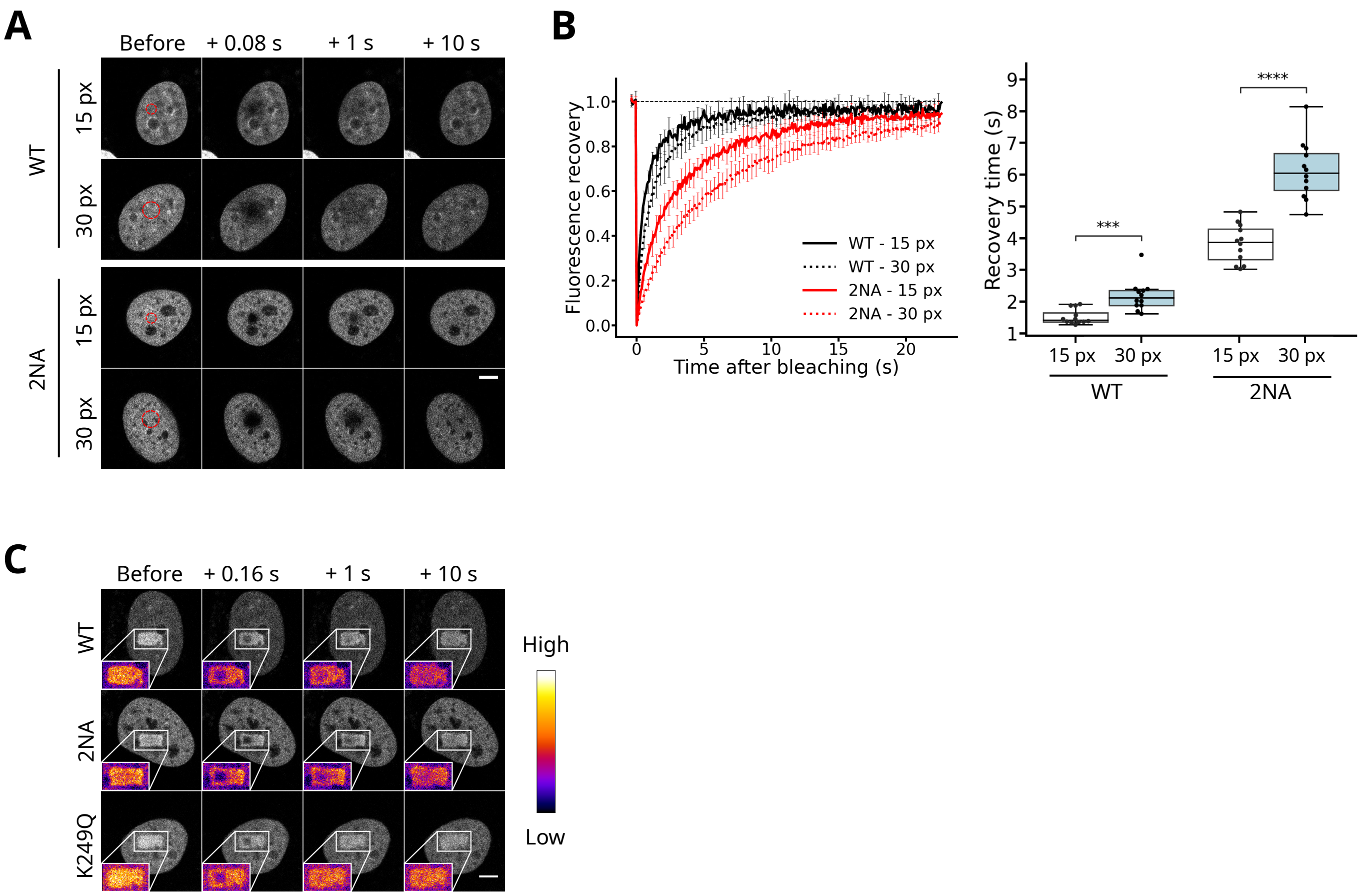


**Figure S5. Nuclear dynamics of the OGG1-N149A/N150A (2NA) mutant in the absence of external damage and at sites of laser micro-irradiation. (A)** Representative time-course images of the fluorescence recovery after photobleaching of circular area of variable sizes within the nucleus of HeLa OGG1 KO cells expressing GFP-tagged OGG1-WT or OGG1-2NA. The bleached regions of 15 and 30 pixel diameters are shown with red dashed circles. Scale bar: 5 µm. **(B)** Left: Normalized fluorescence recovery curves for GFP-tagged OGG1-WT and OGG1-2NA obtained from the images shown in A. Right: Characteristic recovery times estimated from the fit of the curves. 12 cells per condition. **(C)** Representative time-course images of the fluorescence recovery after photobleaching a sub-region of the area subjected to laser micro-irradiation within the nucleus of HeLa OGG1 KO cells expressing GFP-tagged OGG1-WT, OGG1-2NA or OGG1-K249Q. Insets in pseudocolor show a magnified view of the micro-irradiated region. Scale bar: 5 µm.
